# Population Stratification at the Phenotypic Variance level and Implication for the Analysis of Whole Genome Sequencing Data from Multiple Studies

**DOI:** 10.1101/2020.03.03.973420

**Authors:** Tamar Sofer, Xiuwen Zheng, Cecelia A. Laurie, Stephanie M. Gogarten, Jennifer A. Brody, Matthew P. Conomos, Joshua C. Bis, Timothy A. Thornton, Adam Szpiro, Jeffrey R. O’Connell, Ethan M. Lange, Yan Gao, L. Adrienne Cupples, Bruce M. Psaty, NHLBI Trans-Omics for Precision Medicine (TOPMed) Consortium, Kenneth M. Rice

**Author notes:** Correspondence: Tamar Sofer.

## Abstract

In modern Whole Genome Sequencing (WGS) epidemiological studies, participant-level data from multiple studies are often pooled and results are obtained from a single analysis. We consider the impact of differential phenotype variances by study, which we term ‘variance stratification’. Unaccounted for, variance stratification can lead to both decreased statistical power, and increased false positives rates, depending on how allele frequencies, sample sizes, and phenotypic variances vary across the studies that are pooled. We describe a WGS-appropriate analysis approach, implemented in freely-available software, which allows study-specific variances and thereby improves performance in practice. We also illustrate the variance stratification problem, its solutions, and a corresponding diagnostic procedure in data from the Trans-Omics for Precision Medicine Whole Genome Sequencing Program (TOPMed), used in association tests for hemoglobin concentrations and BMI.

---

Large-scale association analyses using whole genome sequence (WGS) data on thousands of participants are now underway, through programs such as the NHLBI’s TOPMed and NHGRI’s Genome Sequencing Program. Unlike earlier Genome-Wide Association Studies (GWASs), where data were combined using meta-analyses of summary statistics, in WGS analyses participant-level data from multiple studies are often pooled, and results are obtained from a single analysis. Pooled analysis of WGS is useful, due to its computational tractability, ability to control for genetic relatedness across the pooled datasets, and its statistical testing procedures’ improved performance when applied to rare variants. However, it is sensitive to a form of population stratification that is not well known. Population stratification in genetic association analysis [1] typically refers to situations where the mean phenotype value and allele frequency both differ across population subgroups. Unless appropriately accounted for in the analysis, e.g. by using regression-based adjustments for ancestry such as principal components or genetic relatedness matrices in linear mixed models, or their combination, it can lead to false-positive associations [2-4]. But population stratification can also manifest as differences in phenotype *variances*, across population subgroups, combined with differences in allele frequencies. In practice, this phenomenon is common in pooled analysis of multi-study data, as small differences in allele frequencies are prevalent, and different studies being pooled often have different measurement protocols, environmental exposures and inclusion criteria, all of which can lead to different phenotype variances among studies.

A standard tool for analysis of quantitative traits is linear or linear mixed model regression. In its widely-used default version, linear regression is fitted under the assumption that the phenotype’s residual variance is the same for all individuals in the analyzed sample. Similarly, linear mixed models are usually fitted under the assumption that the variance components due to environmental effects and measurement error are equally-sized across all studies. The extent of the consequences if the variances are not equal-sized can be computed exactly given simplifying assumptions (see Online Methods). Broadly, using default approaches, if a specific subgroup has a larger phenotypic variance than that of other subgroups in the pooled analysis, the estimated precision of the association signal will understate the contribution from such a subgroup. The result is deflation (loss of power) for variants where allele frequency is greater in this subgroup compared to other subgroups, or inflation (too many false positives) for variants with lower allele frequency in this subgroup compared to others.

While default linear regression methods assume the same variance for all subgroup, which leads to mis-calibrated tests if the assumption does not hold, standard computational tools can be adapted to allow for a stratified variance model, yielding correctly calibrated tests. Specifically, by fitting different residual variances for each study, or more generally, appropriately-defined “analysis group” (e.g. all African Americans of a specific study) the problem can be alleviated. This can be viewed as fitting a different variance component for noise within each study, or as a weighted least squares approach, in which the group-dependent weights are estimated. This approach is implemented in some standard genetic analysis software packages (e.g. GENESIS [5]). Full details, including coding examples, are given in the Online Methods, but here we note that such software exists, and is fast enough for high-throughput analysis in large samples.

We demonstrate the variance stratification problem in analyses of hemoglobin concentrations (HGB, N=7,596; 3 analysis groups) and body mass index (BMI, N=9,807; 8 analysis groups) in the TOPMed freeze 4 dataset. For both traits, we performed single-variant association analysis for all variants with minor allele count of at least 20. Detailed breakdown of the studies and populations used in these analyses are provided in the Online Methods, Tables 1 and 2. The analysis strategy for both traits was: first fit a mixed linear regression model, with fixed effects for sex, age (also age^2^ for BMI), group defined by study and race/ethnicity, and allowing for genetic relatedness by including a variance component proportional to a Genetic Relationship Matrix (GRM [6]) computed on all available variants with minor allele frequency at least 0.001. Then we took the residuals generated by this model, rank-normalized them, and then re-fit the same model but with the rank-normalized residuals as the trait [7]. For both traits, we compared three analyses: first, a ‘homogeneous variance’ analysis that estimates a single residual variance parameter across all individuals; second, a ‘stratified residual variance’ model that allows a different residual variance parameter for each analysis group, and a ‘completely stratified’ approach which fits models and performs tests in each analysis groups separately, and then combines the results via inverse-variance fixed-effects meta-analysis. Analysis groups were either all individuals from a single study (e.g. Amish), or further defined by both study and race/ethnic group (e.g. European and African Americans from the Cleveland Family Study were separate analysis groups). For BMI, we removed 8 individuals from the ‘completely stratified’ analysis to ensure individuals were unrelated across groups, defined as less than third-degree relatedness.

**Table 1:** Analysis groups/strata participating in the BMI analysis. For each group, we report parent study, race/ethnic group, number of participants, number and percentage of males, and age and BMI’s means and standard deviations.

| Study | Race/ethnic group | N | Male sex<br>Number (%) | Age<br>Mean (SD) | BMI<br>Mean(SD) |
| --- | --- | --- | --- | --- | --- |
| JHS | African American | 1756 | 641 (36.5%) | 40.3 (9.9) | 25.8 (4.7) |
| CFS | European American | 471 | 235 (49.9%) | 44.0 (19.6) | 31.0 (9) |
| FHS | European American | 3576 | 1584 (44.3%) | 37.2 (9) | 25.6 (4.8) |
| COPDGene | African American | 881 | 519 (58.9%) | 58.7 (6.7) | 28.4 (6.7) |
| COPDGene | European American | 995 | 505 (50.8%) | 63.8 (8) | 27.9 (5.7) |
| CRA | Costa Rican | 550 | 285 (51.8%) | 18.9 (16.1) | 20.8 (5.3) |
| Amish | European American | 1106 | 556 (50.3%) | 51.9 (16.9) | 27.1 (4.7) |
| CFS | African American | 472 | 205 (43.4%) | 40.0 (19) | 32.7 (9.6) |

**Table 2:** Analysis groups/strata participating in the analysis of hemoglobin concentration. For each group, we report parent study, race/ethnic group, number of participants, number and percentage of males, and age and hemoglobin’s means and standard deviations.

| study | Race/ethnic group | N | Male sex<br>Number (%) | Age<br>Mean (SD) | Hemoglobin<br>Mean (SD) |
| --- | --- | --- | --- | --- | --- |
| JHS | African American | 3251 | 1212 (37.3%) | 54.8 (12.8) | 13.0 (1.5) |
| FHS | European American | 3133 | 1512 (48.3%) | 58.5 (15.0) | 14.1 (1.3) |
| Amish | European American | 1102 | 557 (50.5%) | 50.6 (16.9) | 13.8 (1.2) |

In the online methods, we provide mathematical derivation and code for computing expected variance-specific inflation factors *λ* _*vs*_ and for demonstrating the variance stratification problem, under simplifying assumptions where only residual variance is modeled. Using these, for each variant in the BMI and HGB analyses we computed an approximate variant-specific inflation factor *λ*_*vs*_. We investigated the inflation/deflation problems resulting from variance stratification, and verified that the patterns of inflation and deflation in the homogeneous variance analysis agree, across the different variants, with those obtained from the formula and code provided in the Online Methods. Figures 1 and 2 provide quantile-quantile (QQ) plots for variants from three broad categories of variants, where theory predicts inflation (*λ* _*vs*_ > 1.03), deflation (*λ* _*vs*_ < 0.97) and “about right” (0.97 < *λ* _*vs*_ < 1.03), and across all variants, for HGB and BMI analyses respectively. The plots overlay the results from the three analyses methods together. While the homogeneous variance model clearly produces inflated and deflated QQ-plots in line with the theoretical expectation, when looking at all tested variants together, this inflation and deflation (i.e. Type I and Type II errors) mask each other, alarmingly. Despite appearances, these problems do not “cancel out”; one creates more Type I errors, one creates more Type II errors, yet the plot of all results may lead investigators to conclude that the analysis is well-calibrated. In contrast, the stratified residual variance model provides better control of Type I errors, as seen in the QQ-plots, with the exception of the bottom left panel in Fig. 2, which provides QQ-plots for the set of variants that are expected to have deflated test statistics under the pooled variance model: here the stratified residual variance model was also somewhat deflated. Fig. 3 provide the genomic control inflation factors *λ* _*gc*_ computed over each of the variant sets provided in the QQ-plots and for each of the traits. The completely stratified model performed better in terms of overall QQ-plots and computed *λ* _*gc*_ values in the two analyses.

**Figure 1:**
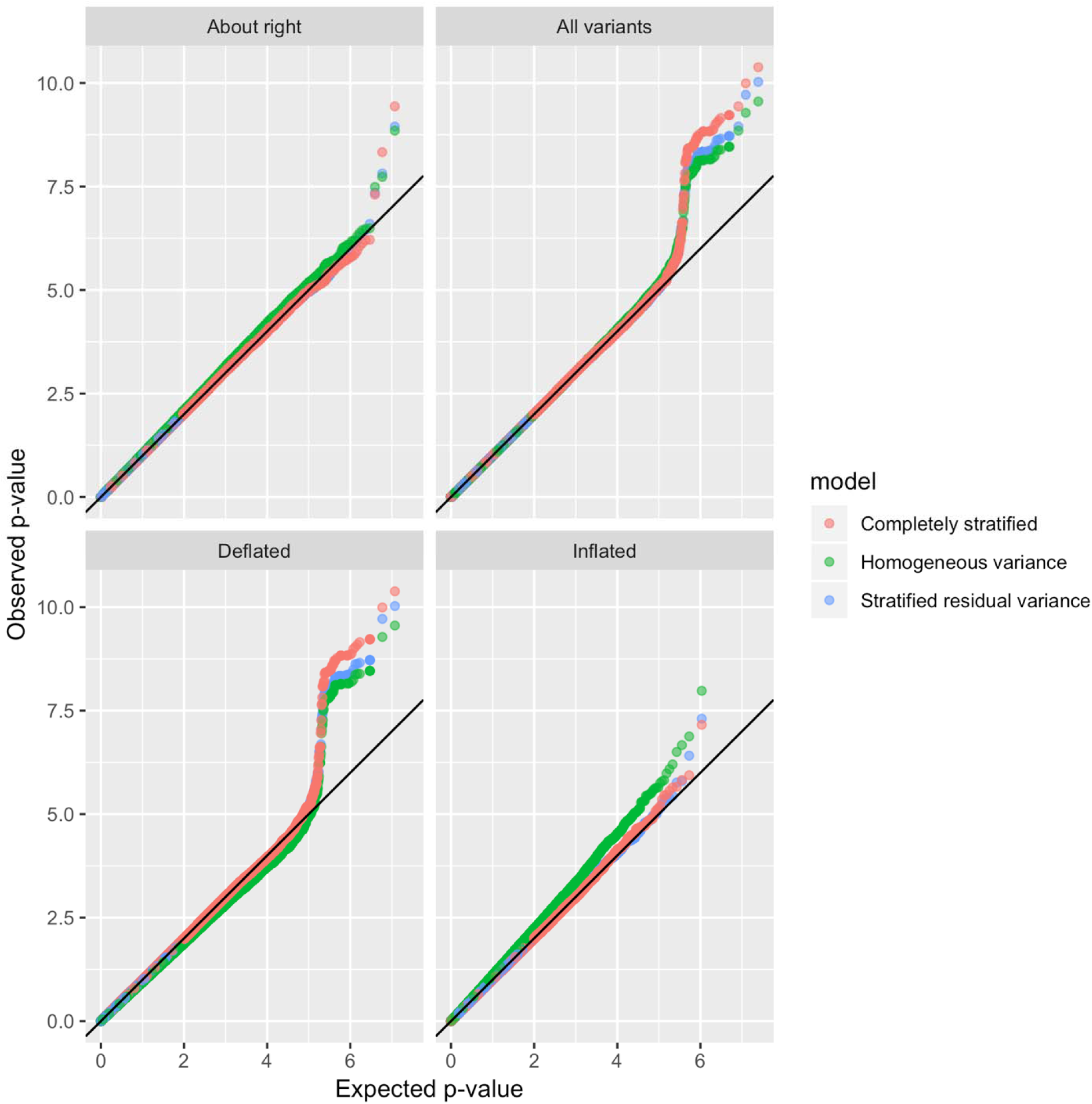
QQ-plots comparing observed and expected p-values from the analysis of hemoglobin concentrations using three approaches: “homogeneous variance” model, that assumes that all groups in the analysis have the same variances; “stratified variance” model, that allows for different residual variances across analysis groups; and a “completely stratified” model in which analysis groups were analyzed separately, allowing for both heterogeneous residual and genetic variances across groups, and then combined together in meta-analysis. The QQ-plots are provided across sets of variants classified by their inflation/deflation patterns according to the algorithm for variant-specific approximate inflation factors. We categorized variants as “About right” when they had estimated between 0.97 to 1.03, “Deflated” when estimated lower than 0.97, and “Inflated” when they had estimated higher than 1.03.

**Figure 2:**
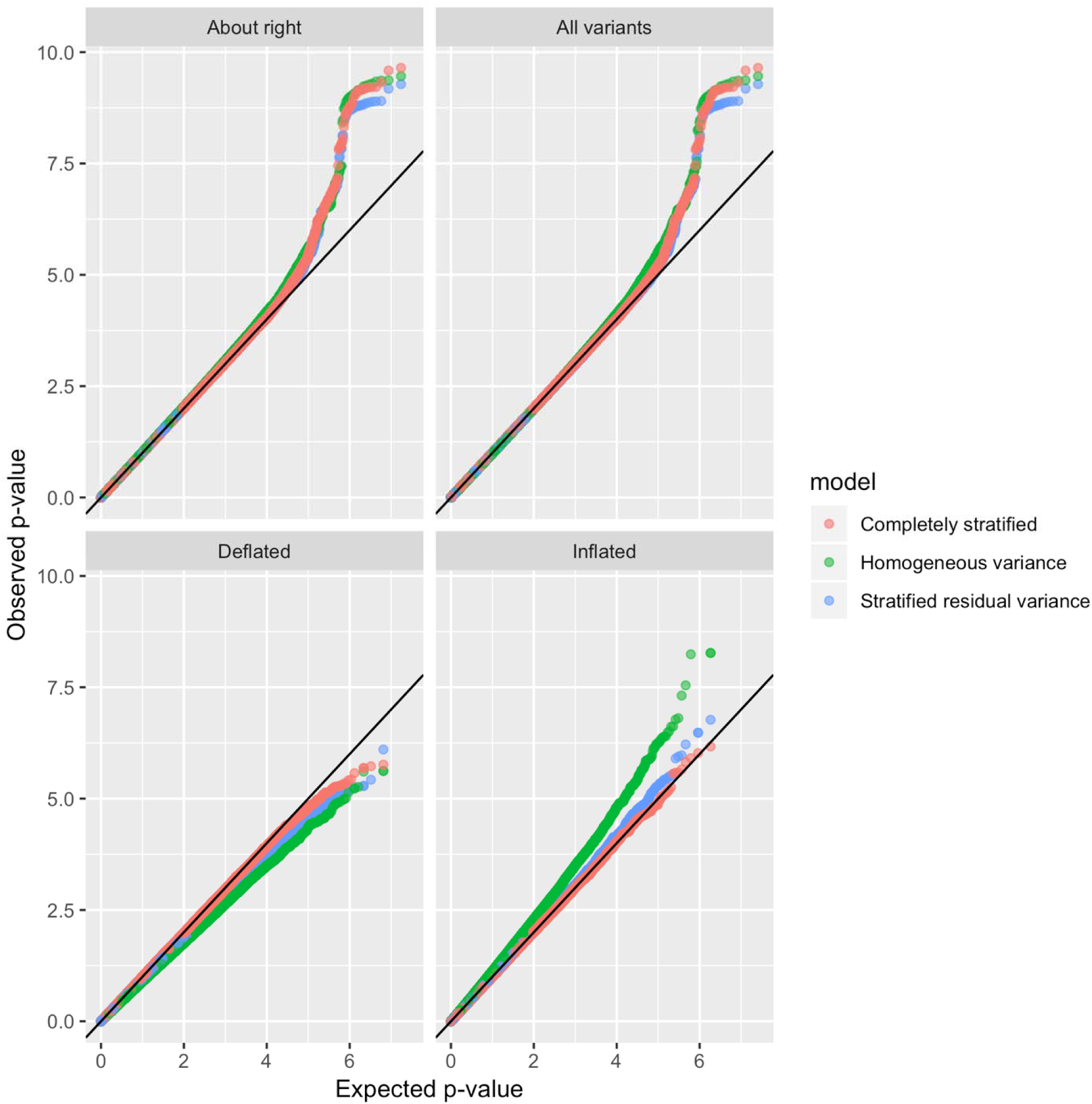
QQ-plots comparing observed and expected p-values from the analysis of BMI using three approaches: “homogeneous variance” model, that assumes that all groups in the analysis have the same variances; “stratified variance” model, that allows for different residual variances across analysis groups; and a “completely stratified” model in which analysis groups were analyzed separately, allowing for both heterogeneous residual and genetic variances across groups, and then combined together in meta-analysis. The QQ-plots are provided across sets of variants classified by their inflation/deflation patterns according to the algorithm for variant-specific approximate inflation factors. We categorized variants as “About right” when they had estimated between 0.97 to 1.03, “Deflated” when estimated lower than 0.97, and “Inflated” when they had estimated higher than 1.03.

**Figure 3:**
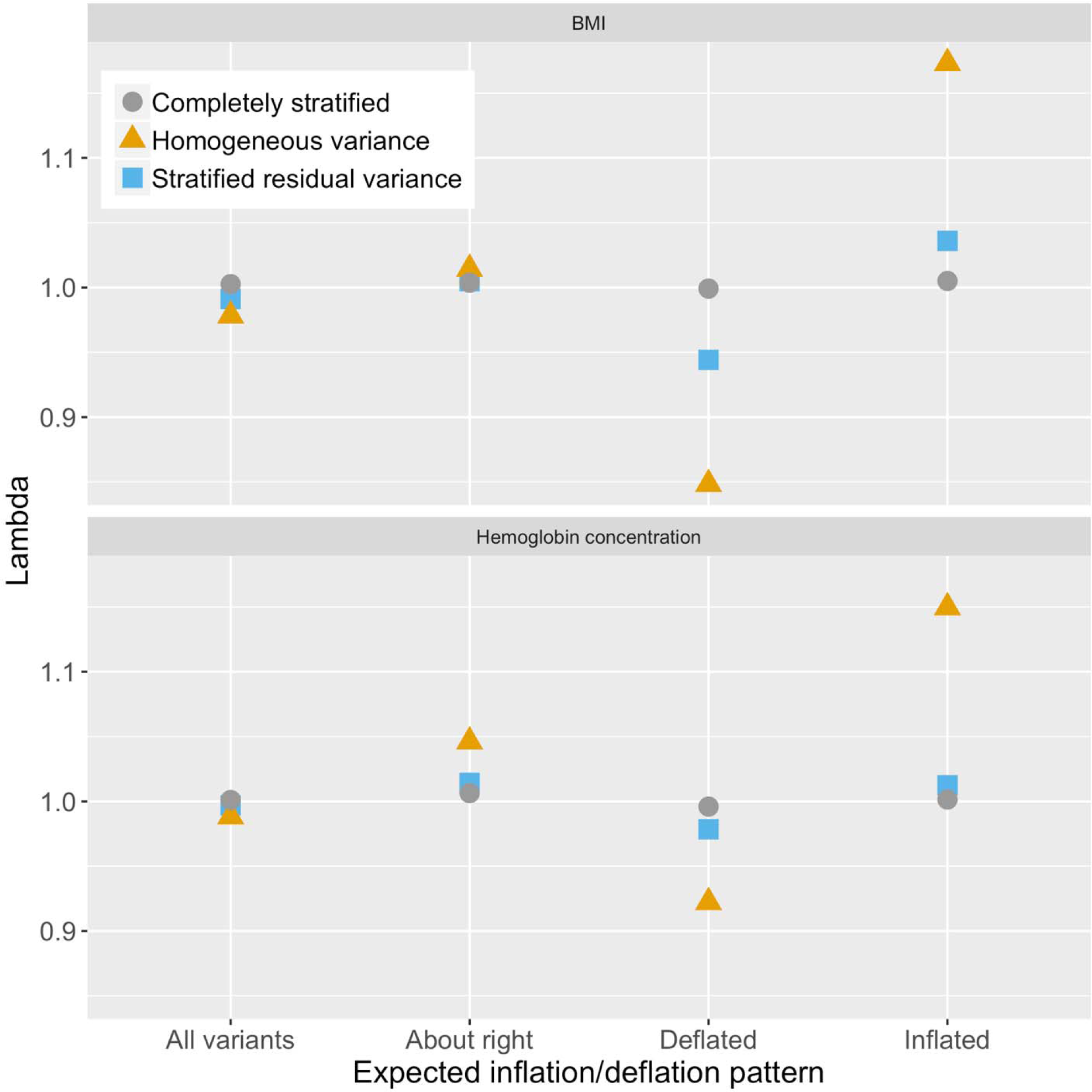
Estimated genomic control inflation factors (in the various analyses of BMI and hemoglobin concentrations, computed across sets of variants classified by their inflation/deflation patterns according to the algorithm for approximate variant-specific inflation factors (. We categorized variants as “About right” when they had estimated between 0.97 to 1.03, “Deflated” when estimated lower than 0.97, and “Inflated” when they had estimated higher than 1.03. Genomic control inflation factors were computed as the ratio between the median test statistic across variants in the set to the theoretical median of the test statistic under the null hypothesis of no association.

In linear regression, the stratified residual variance model allows every analysis group to have its own residual variance parameter. In the mixed model setting, where the variance is decomposed into genetic and residual variance, this model keeps the genetic variance component the same but allows for the residual variance to differ across groups. Analysis groups can be defined as study, race, and combinations of these. Our mathematical derivation and code for computing AVS are under simplifying assumptions of no covariate effects and independent observations. Therefore, these make no distinction between genetic and residual variance components. While in the linear regression setting (independent observations) variance stratification model clearly suffices to account for variance heterogeneity, in the mixed model setting, a residual variance stratification model may not be optimal, because it does not account for stratification in the genetic variance as well, which could be the result of study design. For example, in Fig. 4, the estimated genetic variance component of the Cleveland Family Study is much higher than those of other studies, and of the residual variance component of the same study, perhaps because study participants were selected from families with Obstructive Sleep Apnea, which is highly associated with obesity. Heterogeneity in genetic variance is addressed in the ‘completely stratified model’, but such a model requires that individuals are independent between different groups (strata). A method that allows for complete stratification of analysis groups, while keeping genetically related individuals across these groups, was previously developed [8] and was shown to have good statistical properties. However, it was only introduced for the Wald test, and not the Score test, which is often used for rare variants association analyses, and it is currently computationally costlier than a pooled analysis because individual level data are used both at the individual analysis group computations, and when computing covariances between effect size estimates of all analysis groups. Computational efficiency is critical when testing the large number of variants observed in WGS studies. As sample sizes of TOPMed grow, pooling together more diverse studies and populations, variance stratification problems may be more severe. Models allowing for pooled analysis with both group-specific residual and genetic variances or robust variance estimates may be needed for better control of Type I errors and increased efficiency.

**Figure 4:**
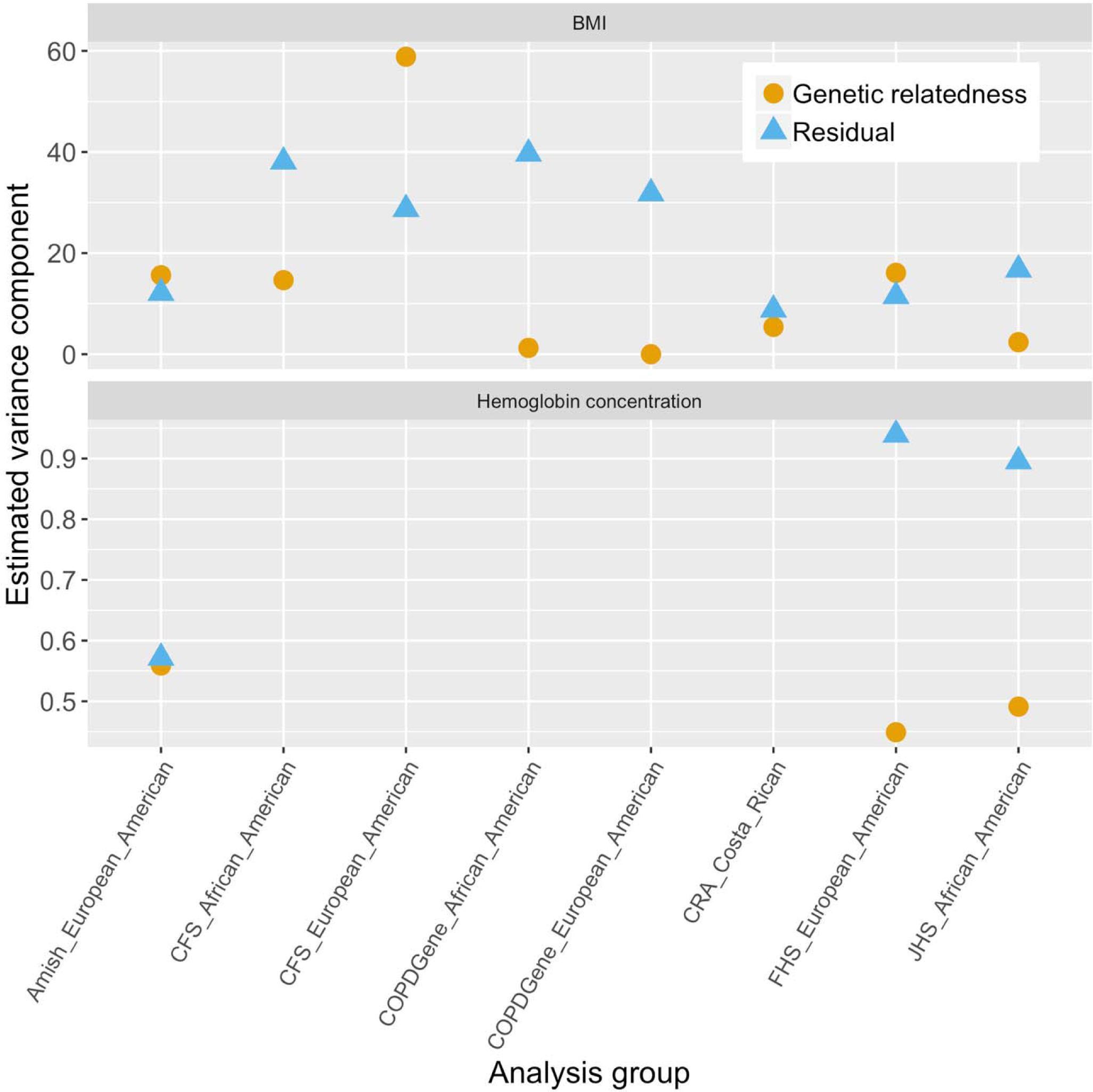
Estimated variance components corresponding to residual and genetic relatedness in the analyses of BMI and hemoglobin concentration. For each analysis group, the estimated variance components were computed based on the analysis of the group alone.

## Supporting information

Tutorial on computing and using variant-specific inflation

## Acknowledgements

T.S. was supported by National Heart, Lung, and Blood Institute (NHLBI; R01HL120393-03S1, 1R35HL135818, and 1R21HL145425). The views expressed in this manuscript are those of the authors and do not necessarily represent the views of the National Heart, Lung, and Blood Institute; the National Institutes of Health; or the U.S. Department of Health and Human Services.

## Author Contribution Statement

K.M.R. and T.S. conceptualized the work and drafted the manuscript. X.Z., S.M.G, M.C., T.A.T., A.S., and K.M.R., developed, studied, and implemented the genetic analysis algorithm that incorporates different residual variances by group. X.Z., C.A.L, S.M.G performed quality control on the genetic sequencing data. X.Z., C.A.L., J.A.B, and J.B., harmonized and performed quality control for the phenotypes used in the analysis. B.M.P, C.C.L, and K.M.R. supervised quality control and phenotype harmonization procedures. All authors interepreted the data, reviewed and approved the final manuscript.

## Online Methods

### Mathematical derivation when observations are independent within study

For a total sample size of *n*, we assume that the data follow a linear model denoted as

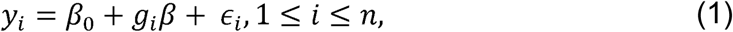

where *Y*_*i*_ is the trait or phenotype value of person *i, g*_*i*_ is their count of coded alleles (i.e. genotype), *β*_0_ denotes the mean outcome in those with no copies of the coded allele, *β* denotes the effect on the mean trait of each additional copy of the coded allele, and the *ϵ*_*i*_ are residual errors, which we for now assume are independent.

To provide intuition for the variance stratification problem, we first demonstrate a mathematical derivation under simplifying assumptions. We assume that the study individuals are independent and identically distributed (iid), that the genotype effect is null (*β* = 0), and that the residuals follow a normal distribution *ϵ*_*i*_ ∼ *N*(0, *σ*^2^). We further assume that the phenotypes are centered, the genotypes are centered, and follow a dominant mode of inheritance, i.e. we are using 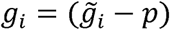, where 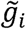 is the genotype under a dominant mode (having values 1 or 0), is its frequency, and *g*_*i*_ is used in analysis.

#### The Wald test

The Wald test quantifies the significance of the data by dividing a regression-based estimate of *β* by its corresponding estimated standard error. The linear regression estimate of the effect is

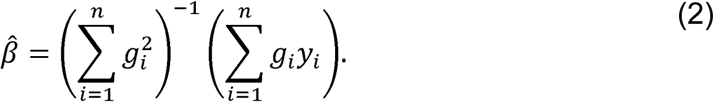

Denoting the residual variance of individual *i* by 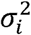 (which may differ across individuals), the variance of 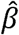 is

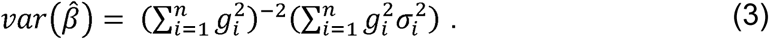

When the variance of the residuals is homogeneous across all individuals, this is

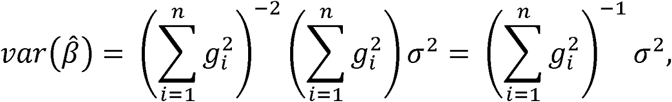

where *σ*^2^ is the common variance. To illustrate how this approach can mislead under heterogeneous variance, we consider the situation where two subgroups are present, of sizes *n*_1_ and *n*_2_ respectively, such that *n*_1_ + *n*_2_ = *n*. Further, each study is internally homogeneous with residual variances 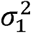 and 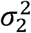, and it is also useful to write *p*_1_ and *p*_2_ for the frequency of the variant of interest (under dominant mode). Because we assume that the variant was centered in the pooled population, we have that 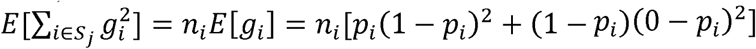, or 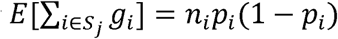.We can re-write equation (3) as:

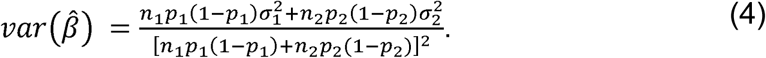

We see that the actual variance is a linear combination of the variance parameters 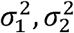, and the weight assigned to each depends on the minor allele frequency and sample size in each group. Where the minor allele frequencies are equal the two forms are equal, as there is no association between genotype and outcome, and no confounding occurs. But where they are not equal then the variance of the estimator upweights the residual variance in the group where the variant is more common, which does not happen under homogeneity. This result straightforwardly generalizes for *M* analysis groups. Next, we provide a more general algorithm for computing variant-specific inflation factors.

### Computing approximate variant-specific inflation factors

We can also use mathematical derivations under homogeneity and heterogeneity, relaxing the restrictive assumptions provided earlier, to compute variant-specific inflation factors, using the standard Huber “sandwich” formula [9]. We now allow for additive genetic model, and do not assume that the genotypes are standardized. The variance estimator used by the Wald test is now provided as follows:

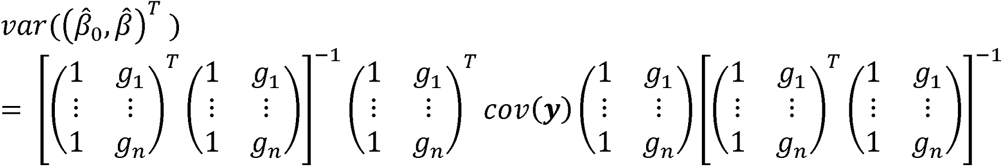

which simplifies if *cov*(*y*_*i*_) = *σ*^2^ for all *i* = 1, …, *n* to:

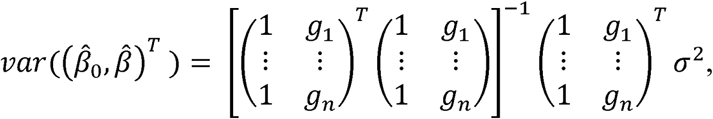

but allowing for different variances per analysis groups, it becomes:

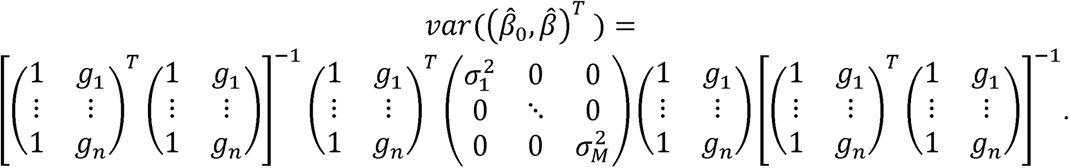

Based on these two expressions, we propose an algorithm to compute an approximate variant-specific inflation factor. For computational purposes, we further simplify these arguments taking advantage of the fact that there are repeated rows (e.g. corresponding to people who have *g*_*i*_ = 1 are from the same analysis unit, i.e. have the same residual variance). The algorithm below uses the additional assumption that phenotype variance within each study does not vary with genotype – which must hold under the strong null hypothesis of no association in any subpopulation. It also uses the simplifying assumption that variants are in Hardy Weinberg Equilibrium (HWE) within each study population; testing HWE is a standard pre-processing step for genotype data.

#### Algorithm for computing variant-specific inflation factors

Define:

- *X* = [*G D*] where *G* is a vector of length *3M* (giving all options of number of counts of an allele in the *M* analysis groups) in which element *i* is 0: if *i* = 3*m* − 2, 1: if *i* = 3*m* − 1, 2: if *i* = 3*m*, for some integer *m* between 1 and *M*, and matrix *D*is a 3*M* × *M* design matrix where the *i, j* element is 1: if *i* = 3*m*, 3*m* − 1 *or* 3*m* − 2 for some integer *m* between 1 and *M* AND *j* = *m* for the same *m*, 0: otherwise
- *P*, a 3*M* × 3*M* diagonal matrix, in which each entry gives the population proportion in each combination of genotype and analysis group.
- *V*, is a 3*M* × 3*M* diagonal matrix, in which each entry gives the outcome variance in each combination of genotype and analysis group.

Under heterogeneity the variance of the slope estimate is proportional to the leading diagonal entry of the “sandwich” form *B*^−1^*AB*^−1^, where *B* = *X*^*T*^ *PX* and matrix *A* = *X*^T^ *PVX*. Under homogeneity the variance of the slope estimate is proportional to the leading entry of *B*^−1^ × *sum*(*diag*(*PV*)), with the same constant of proportionality. The ratio of these two leading entries gives the large-sample value of *λ*_*gc*_, the genomic control inflation factor [10] that would be obtained by comparing the median Wald test statistic to the median of the 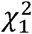 reference distribution. Because this formula provides different results for each variant, depending on the allele frequencies, we denote the ratio between the estimated values under homogeneous variance and the heterogeneous variance models *λ*_*vs*_, for “variant specific”.

An R function implementing these matrix calculations is provided, together with a tutorial including a coding example. These are also provided on GitHub on https://github.com/tamartsi/Variant_specific_inflation.

### Whole Genome Sequencing in TOPMed

Whole genome sequencing (WGS) was performed on DNA samples extracted from blood. Sequencing was performed by the Broad Institute of MIT and Harvard (FHS and Amish) and by the Northwest Genome Center (JHS). PCR-free libraries were constructed using commercially available kits from KAPA Biosystems (Broad) or Illumina TruSeq (NWGC). Libraries were pooled for clustering and sequencing, and later de-multiplexed using barcodes. Cluster amplification and sequencing were performed according to manufacturer’s protocols using the Illumina cBot and HiSeq X sequencer, to a read depth of >30X. Base calling was performed using Illumina’s Real Time Analysis 2 (RTA2) software. Read alignment, variant detection, genotype calling and variant filtering were performed by the TOPMed Informatics Research Center (University of Michigan). Reads were aligned to the 1000 Genomes hs37d5 decoy reference sequence. Variant detection and genotype calling were performed jointly for several TOPMed studies (including the three analyzed here), using the GotCloud pipeline. Mendelian consistency was used to train a variant quality classifier using a Support Vector Machine, used for variant filtering. Additional quality control (pedigree checks, gender checks, and concordance with prior array data), performed by the TOPMed Data Coordinating Center, were used to detect and resolve sample identity issues. Further details (including software versions) are provided online (see URL).

### URL

https://www.ncbi.nlm.nih.gov/projects/gap/cgi-bin/document.cgi?study_id=phs000964.v2.p1&phv=251960&phd=6969&pha=&pht=4838&phvf=&phdf=&phaf=&phtf=&dssp=1&consent=&temp=1

### TOPMed studies in the analyses

#### Participant characteristics: BMI analysis

#### Framingham Heart Study

The Framingham Heart Study (FHS) acknowledges the support of contracts NO1-HC-25195 and HHSN268201500001I from the National Heart, Lung and Blood Institute and grant supplement R01 HL092577-06S1 for this research. We also acknowledge the dedication of the FHS study participants without whom this research would not be possible. FHS participated in both the hemoglobin and BMI analysis.

#### Jackson Heart Study

JHS is a longitudinal community-based study designed to assess the causes of the high prevalence of cardiovascular disease among AAs in the Jackson, Mississippi metropolitan area [11]. During the baseline examination period (2000-2004) 5,306 self-identified AAs were recruited from urban and rural areas of the three counties (Hinds, Madison and Rankin) that comprise the Jackson, Mississippi metropolitan area. Participants were between 35 and 84 years old with the exception of a nested family cohort, where those ≥21 years old were eligible All participants included in analyses provided written informed consent for genetic studies. Approval was obtained from the institutional review board of the University of Mississippi Medical Center (UMMC). Data on participants’ health behaviors, medical history, and medication use were collected at baseline and subjects underwent venipuncture, allowing for assessment of complete blood cell counts and other measures at UMMC (Beckman-Coulter, [12]). JHS participated in both the hemoglobin and BMI analysis.

The Jackson Heart Study (JHS) is supported and conducted in collaboration with Jackson State University (HHSN268201300049C and HHSN268201300050C), Tougaloo College (HHSN268201300048C), and the University of Mississippi Medical Center (HHSN268201300046C and HHSN268201300047C) contracts from the National Heart, Lung, and Blood Institute (NHLBI) and the National Institute for Minority Health and Health Disparities (NIMHD). The authors also wish to thank the staffs and participants of the JHS.

#### The Amish study

We gratefully acknowledge our Amish liaisons, research volunteers, field workers and Amish Research Clinic staff and the extraordinary cooperation and support of the Amish community without which these studies would not have been possible. The Amish studies are supported by grants and contracts from the NIH, including U01 HL072515, U01 HL84756, U01 HL137181 and P30 DK72488. The Amish study participated in both the hemoglobin and BMI analysis.

#### The Genetic Epidemiology of Asthma in Costa Rica – Asthma Costa Rica Cohort

Costa Rica is a Hispanic population with asthma prevalence 24%. This study originated in 1997 with the recruitment of both extended pedigrees and trios in the central valley of Costa Rica. The study participated in the BMI analysis.

#### Cleveland Family Study

The Cleveland Family Study (CFS, [13]) was designed to examine the genetic basis of sleep apnea in 2,534 African-American and European-American individuals from 356 families. CFS participated in the BMI analysis.

#### COPDGene

COPDGene is a cross sectional prospective cohort enrolled between January 2008 and June 2011 at 21 Clinical Centers. The study goals were to characterize smokers with and without COPD, using spirometry, six minute walk, medical history, respiratory symptoms (modified ATS respiratory questionnaire), respiratory medications, quality of life and inspiratory and expiratory chest CT scans, and to perform epidemiological and genetic studies of these COPD-related phenotypes. Participants from COPDGene who consented for data use for method development participated in the BMI analysis.

The COPDGene project was supported by Award Number U01 HL089897 and Award Number U01 HL089856 from the National Heart, Lung, and Blood Institute. The content is solely the responsibility of the authors and does not necessarily represent the official views of the National Heart, Lung, and Blood Institute or the National Institutes of Health. The COPDGene project is also supported by the COPD Foundation through contributions made to an Industry Advisory Board comprised of AstraZeneca, Boehringer Ingelheim, GlaxoSmithKline, Novartis, Pfizer, Siemens and Sunovion.

## TOPMed acknowledgements

Whole genome sequencing (WGS) for the Trans-Omics in Precision Medicine (TOPMed) program was supported by the National Heart, Lung and Blood Institute (NHLBI). WGS for “NHLBI TOPMed: Genetics of Cardiometabolic Health in the Amish” (phs000956) was performed at the Broad Institute of MIT and Harvard (3R01HL121007-01S1). WGS for “NHLBI TOPMed: Cleveland Family Study - WGS Collaboration” (phs000954) was performed at the University of Washington Northwest Genomics Center (3R01HL098433-05S1, HHSN268201600032I). WGS for “NHLBI TOPMed: Genetic Epidemiology of COPD Study” (phs000951) was performed at the University of Washington Northwest Genomics Center (3R01HL089856-08S1), and at the Broad Institute of MIT and Harvard (HHSN268201500014C). WGS for “NHLBI TOPMed: The Genetic Epidemiology of Asthma in Costa Rica - Asthma in Costa Rica cohort” (phs000988) was performed at the University of Washington Northwest Genomics Center (3R37HL066289-13S1, HHSN268201600032I). WGS for “NHLBI TOPMed: Framingham Heart Study” (phs000974) was performed at the Broad Institute of MIT and Harvard (3R01HL092577-06S1, 3U54HG003067-12S2). WGS for “NHLBI TOPMed: Jackson Heart Study” (phs000964) was performed at the University of Washington Northwest Genomics Center (HHSN268201100037C).

Centralized read mapping and genotype calling, along with variant quality metrics and filtering were provided by the TOPMed Informatics Research Center (3R01HL-117626-02S1; contract HHSN268201800002I). Phenotype harmonization, data management, sample-identity QC, and general study coordination were provided by the TOPMed Data Coordinating Center (3R01HL-120393-02S1; contract HHSN268201800001I). We gratefully acknowledge the studies and participants who provided biological samples and data for TOPMed.

