## Supplementary material for "Population Stratification at the Phenotypic Variance level and Implication for the Analysis of Whole Genome Sequencing Data from Multiple Studies": Tutorial on computing and using variant-specific inflation

### Variant-specific inflation factors: a tutorial

Tamar Sofer et al.

2020-03-03

#### Contents

**About this tutorial** **1**

**Preparing for analysis based on simulated data** **2**

**Computing approximate inflation factors based on allele frequencies, sample sizes, and residual standard deviations** **5**

**Performing associating testing: homogeneous variance model** **6**

**Making a QQ-plot figure by categories of inflation factors: only homogeneous variance model** **7**

**Performing associating testing: heterogeneos variance model** **8**

**Making a QQ-plot figure by categories of inflation factors: both homogeneous and stratified variance models** **9**

**Final note** **10**

**Record R package versions** **10**

### About this tutorial

This tutorial demonstrates how to use the R functions provided with the manuscript "Population Stratification at the Phenotypic Variance level and Implication for the Analysis of Whole Genome Sequencing Data from Multiple Studies" to investigate potential mis-calibration of test statistics when testing genetic variant associations with a quantitative trait. P-values may be miscalibrated when individual-level data from multiple studies, or when multiple race/ethnic groups are pooled together, and both allele frequencies and trait residual variances differ between the groups.

In what follows, we will use data that were simulated in advance to compute residual variances and variant allele frequencies, and to test variant associations. Then, we will use this information to generate a figure with multiple panels of QQ-plots, each demonstrating inflation patterns in a different set of genetic variants.

Note: this tutorial uses genetic data saved on a GDS file, and therefore, uses a specific set of R functions to perform tasks such as association testing. You can use other tools and skip to the point where you already have allele frequencies and p-values computed, even if they were computed in another software.

#### Preparing for analysis based on simulated data

In the following code we load packages, "source" functions used, and load the simulated data.

Clean up the workspace, load required packages:

```r
rm(list = ls())
setwd("~/Documents/GitHub/Variant_specific_inflation")
source("variant_specific_inflation_functions.R")
```

```
## Loading required package: tidyr

## Loading required package: data.table

## data.table 1.12.8 using 4 threads (see ?getDTthreads).  Latest news: r-datatable.com

## Loading required package: ggplot2
```

```r
require(GWASTools)
```

```
## Loading required package: GWASTools

## Loading required package: Biobase

## Loading required package: BiocGenerics

## Loading required package: parallel

##
## Attaching package: 'BiocGenerics'
```

```
## The following objects are masked from 'package:parallel':
##
##     clusterApply, clusterApplyLB, clusterCall, clusterEvalQ, clusterExport, clusterMap, parApply, pa[illegible]
##     parSapply, parSapplyLB

## The following objects are masked from 'package:stats':
##
##     IQR, mad, sd, var, xtabs

## The following objects are masked from 'package:base':
##
##     anyDuplicated, append, as.data.frame, basename, cbind, colnames, dirname, do.call, duplicated, e[illegible]
##     intersect, is.unsorted, lapply, Map, mapply, match, mget, order, paste, pmax, pmax.int, pmin, pm[illegible]
##     rownames, sapply, setdiff, sort, table, tapply, union, unique, unsplit, which, which.max, which.[illegible]

## Welcome to Bioconductor
##
##     Vignettes contain introductory material; view with 'browseVignettes()'. To cite Bioconductor, se[illegible]
##     'citation("pkgname")'.
```

```r
require(dummies)
```

```
## Loading required package: dummies

## dummies-1.5.6 provided by Decision Patterns
```

```r
require(data.table)
require(GENESIS)
```

```
## Loading required package: GENESIS
```

Read genotype data, stored in a SNP-GDS file:

```r
gds <- GdsGenotypeReader("Genotypes.gds")
scanID <- getScanID(gds)
snpID <- getSnpID(gds)
geno <- getGenotype(gds)
rownames(geno) <- snpID
colnames(geno) <- scanID
```

```r
close(gds)
```

Load the phenotype data. The data was simulated and stored in a data.frame which we will call `phen`. The quantitative trait we will use is called `trait`. It the simulations, we assume that there are three different groups, `g1, g2, g3`, each has different error standard deviation. Assuming that in genetic analysis we will only adjust for age, we regress the trait on age and take the residuals to compute residual standard deviation.

```r
# load the simulated data:
phen <- getobj("Phenotypes.RData")
phen <- data.table(phen)

# this is how the data looks like:
phen
```

```
##      scanID group      age    trait
##   1:     p1    g1 37.33547 49.14920
##   2:     p2    g1 46.20567 63.54733
##   3:     p3    g1 38.15073 51.74716
##   4:     p4    g1 41.85987 58.10859
##   5:     p5    g1 38.08240 52.27407
##  ---
## 496:   p496    g3 46.12091 71.57149
## 497:   p497    g3 35.44966 51.16452
## 498:   p498    g3 45.51238 65.47968
## 499:   p499    g3 43.73817 64.97567
## 500:   p500    g3 41.66499 54.46750
```

```r
# compute residual standard deviations by group after regressing on age:
residual_SDs <- phen[, sd(lm(trait ~ age)$resid), by = "group"]
residual_SDs_vec <- residual_SDs$V1
names(residual_SDs_vec) <- residual_SDs$group

# these are the estimated residual standard deviations:
residual_SDs_vec
```

```
##       g1       g2       g3
## 1.074580 1.867516 3.193658
```

The approximate inflation factors will also use the sample sizes of the different groups that were pooled together:

```r
# compute group sample sizes
ns <- table(phen$group)

# make sure that the order matches that of the residual SDs vector:
ns <- ns[names(residual_SDs_vec)]
```

We are going to perform association testing with the genotypes. We will now compute allele frequencies across the genotypes that will be tested. These frequencies will be used when computing inflation factors. In the following code, we compute allele frequencies by group. One could use built-in functions, but here is a code that assumes genotypes are on autosomes and computes the frequencies:

```r
freq_list <- vector(mode = "list", length = length(unique(phen$group)))
names(freq_list) <- unique(phen$group)
for (i in 1:length(freq_list)){
    freq_list[[i]] <- rowMeans(geno[,phen[group == names(freq_list[i])]$scanID],
                                                                    na.rm = TRUE)/2
}

# put the frequencies in a data.frame:
eafs <- do.call(cbind, freq_list)

# make sure that the columns order matches the SD vector:
eafs <- eafs[,names(residual_SDs_vec)]
```

#### Computing approximate inflation factors based on allele frequencies, sample sizes, and residual standard deviations

Here we use the function `compute_variant_inflation_factor` which we provide.

```r
expected_inf <- rep(NA, nrow(eafs))
for (j in 1:length(expected_inf)){
        cur_eafs <- eafs[j,]
        if (sd(cur_eafs) == 0) next
        expected_inf[j] <-
          compute_variant_inflation_factor(eafs = cur_eafs,
                                           ns = ns,
                                           sigmas = residual_SDs_vec)[["Inflation_factor"]]
}

exp_inf <- data.frame(variant_id = snpID,
                      expected_inf = expected_inf,
                      stringsAsFactors = FALSE)
```

#### Performing associating testing: homogeneous variance model

We use the R/Bioconductor GENESIS package. We first fit a "null model", and then use it for association testing. In the code, we fit the null model twice, because we use the fully-adjusted two stage procedure described in Sofer et al. (2019, Gen Epi).

```r
phen$group <- as.factor(phen$group)
phen <- data.frame(phen)
# fit null model (homogeneous residual variance).
nullmod_homogeneous <- fitNullModel(phen,
                                    outcome = "trait",
                                    covars = c("group","age"),
                                    verbose = FALSE)
nullmod_homogeneous_rn <- nullModelInvNorm(nullmod_homogeneous,
                                           norm.option = "all",
                                           rescale = "residSD",
                                           verbose = FALSE)
```

```r
### association tests
gds <- GdsGenotypeReader("Genotypes.gds")
genoData <- GenotypeData(gds, scanAnnot = ScanAnnotationDataFrame(phen))
iterator <- GenotypeBlockIterator(genoData)
assoc_homogeneous <- assocTestSingle(iterator, nullmod_homogeneous_rn)
close(gds)
```

#### Making a QQ-plot figure by categories of inflation factors: only homogeneous variance model

We use the function `qq_plot_by_region` to visualize inflation in sets of genetic variants. We use the individual inflation factors that we compued. To define the values for categorizing a variant as "inflated" we used 1.03 and higher; we set the value for which we categories a variant as "deflated" to 0.97 and lower. Finally, we provide a file name for the figure to print to.

Note that you can include results from a few different models (i.e. more than two models) according to the number of columns in the R `data.frame` that contain p-values.

```r
pval_df <- data.frame(variant_id = assoc_homogeneous$variant.id,
                        homogeneous_variance = assoc_homogeneous$Score.pval)
exp_inf <- exp_inf[match(pval_df$variant_id, exp_inf$variant_id),]

qq_plot_by_region(pval_df = pval_df[,c("homogeneous_variance"), drop = FALSE],
                    expected_inf = exp_inf$expected_inf,
                    inflated_val = 1.03,
                    deflated_val = 0.97,
                    thin_high_pvals = 1e5,
                    figure_file_name = "qq_plots_homo_var.png")
```

```
## Saving 6.5 x 4.5 in image
```

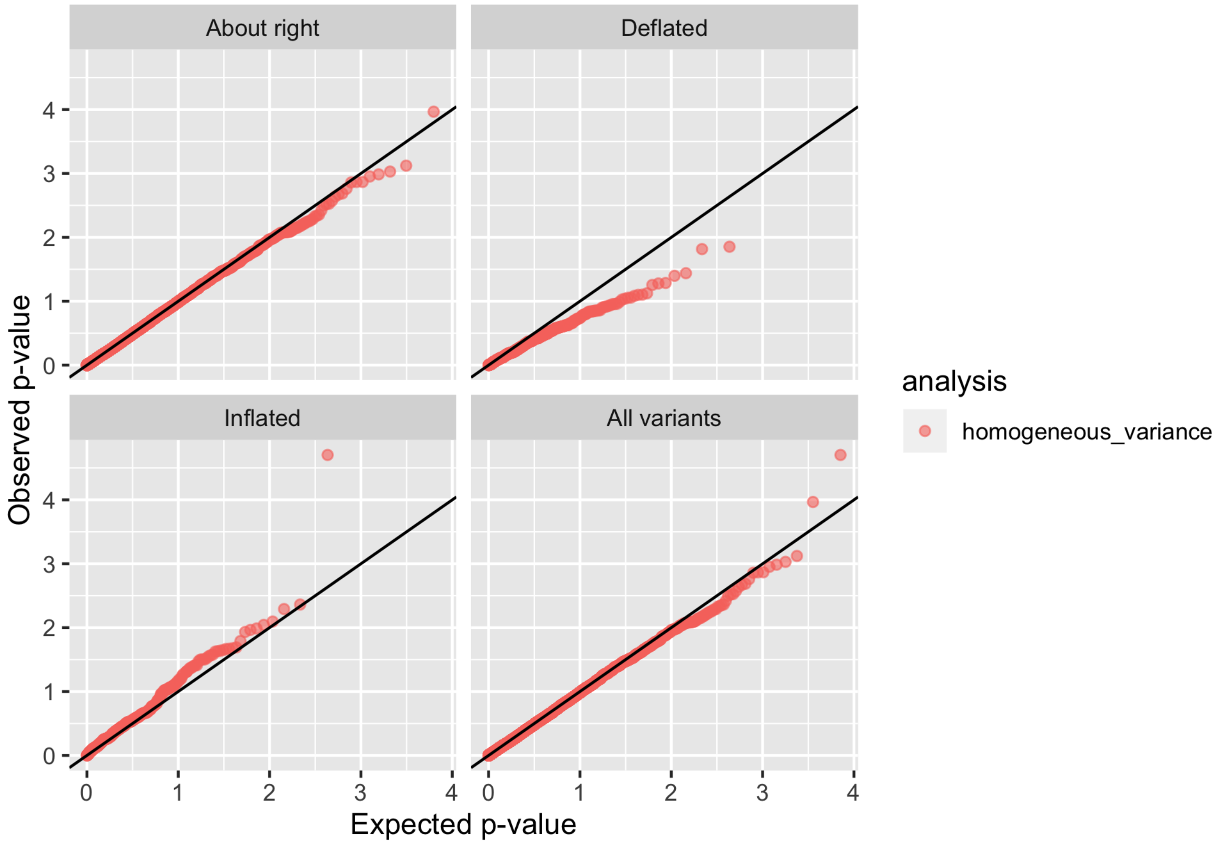


#### Performing associating testing: heterogeneos variance model

```
nullmod_hetero <- fitNullModel(phen,
                               outcome = "trait",
                               covars = c("group","age"),
                               group.var = "group",
                               verbose = FALSE)
nullmod_hetero_rn <- nullModelInvNorm(nullmod_hetero,
                                      norm.option = "all",
                                      rescale = "residSD",
                                      verbose = FALSE)


# testing again:
gds <- GdsGenotypeReader("Genotypes.gds")
genoData <- GenotypeData(gds, scanAnnot = ScanAnnotationDataFrame(phen))
```

```r
iterator <- GenotypeBlockIterator(genoData)
assoc_hetero <- assocTestSingle(iterator, nullmod_hetero_rn)
close(gds)
```

#### Making a QQ-plot figure by categories of inflation factors: both homogeneous and stratified variance models

```r
pval_df <- data.frame(variant_id = assoc_homogeneous$variant.id,
                      homogeneous_variance = assoc_homogeneous$Score.pval,
                          stratified_variance = assoc_hetero$Score.pval)

exp_inf <- exp_inf[match(pval_df$variant_id, exp_inf$variant_id),]

qq_plot_by_region(pval_df = pval_df[,c("homogeneous_variance","stratified_variance")],
                  expected_inf = exp_inf$expected_inf,
                  inflated_val = 1.03,
                  deflated_val = 0.97,
                  thin_high_pvals = 1e5,
                  figure_file_name = "qq_plots_homo_strat_var.png")
```

```
## Saving 6.5 x 4.5 in image
```

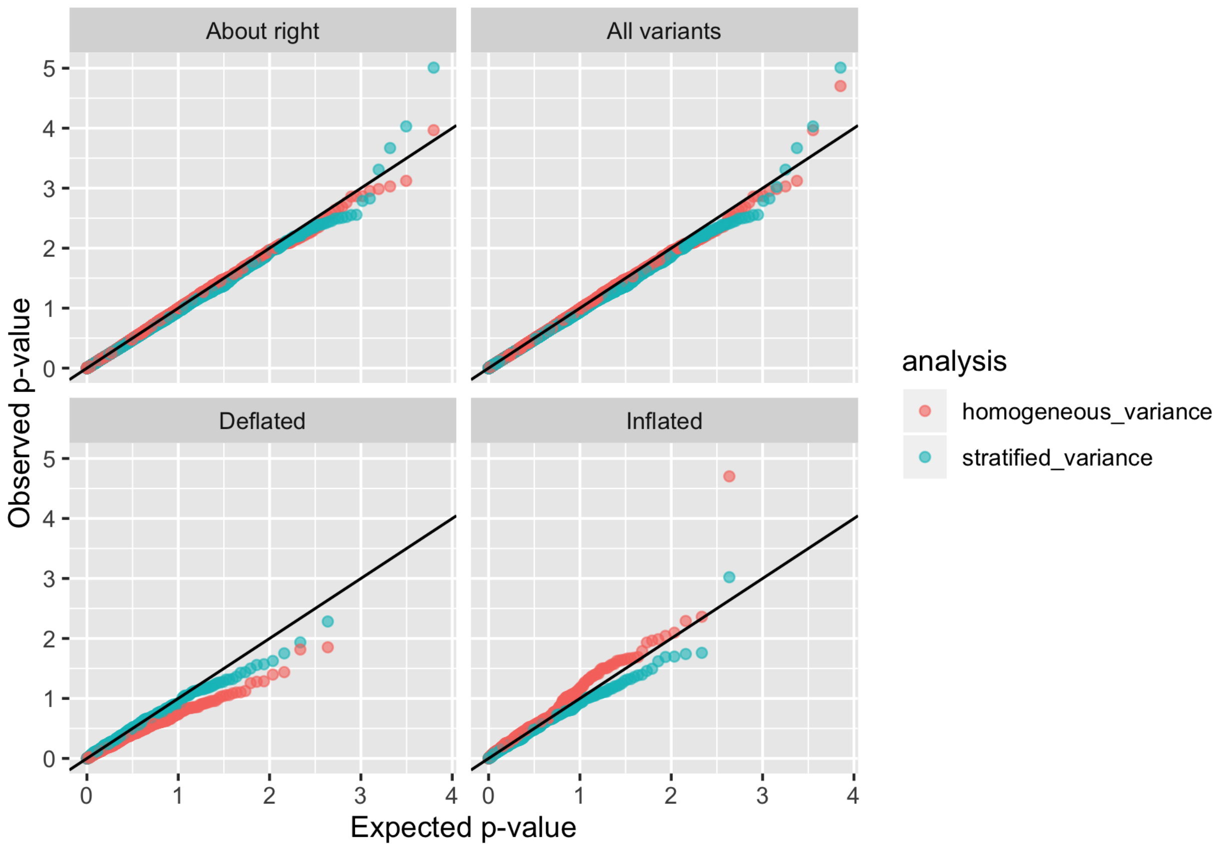


#### Final note

The figures here may not be very impressive. This is because we used a very small dataset, with a small number of people and a small number of variants. For the stratified variance model, this is a small number of people to estimate the group-specific variances. This is likely what causes the perhaps deflation pattern seen in the stratified variance model in the "inflated" variants category.

#### Record R package versions

```r
sessionInfo()
```

```
## R version 3.6.2 (2019-12-12)
## Platform: x86_64-apple-darwin15.6.0 (64-bit)
## Running under: macOS Catalina 10.15.3
##
## Matrix products: default
```

```
## BLAS:   /System/Library/Frameworks/Accelerate.framework/Versions/A/Frameworks/vecLib.framework/Versi
## LAPACK: /Library/Frameworks/R.framework/Versions/3.6/Resources/lib/libRlapack.dylib
##
## locale:
## [1] en_US.UTF-8/en_US.UTF-8/en_US.UTF-8/C/en_US.UTF-8/en_US.UTF-8
##
## attached base packages:
## [1] parallel  stats     graphics  grDevices utils     datasets  methods   base
##
## other attached packages:
## [1] GENESIS_2.16.1      dummies_1.5.6       GWASTools_1.32.0    Biobase_2.46.0      BiocGenerics_0.3
## [8] tidyr_1.0.0         rmarkdown_2.1
##
## loaded via a namespace (and not attached):
##  [1] nlme_3.1-143           bitops_1.0-6           bit64_0.9-7            GenomeInfoDb_1.22.0    too
##  [7] R6_2.4.1               rpart_4.1-15           DBI_1.1.0              lazyeval_0.2.2         mgc
## [13] jomo_2.6-10            SNPRelate_1.20.1       nnet_7.3-12            DNAcopy_1.60.0         wit
## [19] bit_1.1-15.1           compiler_3.6.2         quantreg_5.54          mice_3.7.0             Spa
## [25] labeling_0.3           scales_1.1.0           lmtest_0.9-37          quantsmooth_1.52.0     str
## [31] minqa_1.2.4            GWASExactHW_1.01       XVector_0.26.0         pkgconfig_2.0.3        htm
## [37] rlang_0.4.2            RSQLite_2.2.0          farver_2.0.3           generics_0.0.2         zoo
## [43] RCurl_1.98-1.1         magrittr_1.5           GenomeInfoDbData_1.2.2 Matrix_1.2-18          Rcp
## [49] S4Vectors_0.24.3       lifecycle_0.1.0        stringi_1.4.5          yaml_2.2.0             MAS
## [55] grid_3.6.2             blob_1.2.1             mitml_0.3-7            crayon_1.3.4           lat
## [61] splines_3.6.2          zeallot_0.1.0          knitr_1.27             pillar_1.4.3           Gen
## [67] logistf_1.23           codetools_0.2-16       gdsfmt_1.22.0          stats4_3.6.2           pan
## [73] evaluate_0.14          vctrs_0.2.1            nloptr_1.2.1           foreach_1.4.7          Mat
## [79] purrr_0.3.3            SeqArray_1.26.2        assertthat_0.2.1       xfun_0.12              bro
## [85] survival_3.1-8         tibble_2.1.3           iterators_1.0.12       memoise_1.1.0          IRa
```
